# Genetic characterization of *Streptococcus pyogenes emm*89 strains isolated in Japan from 2011 to 2019

**DOI:** 10.1101/2020.04.14.042051

**Authors:** Yujiro Hirose, Masaya Yamaguchi, Norihiko Takemoto, Tohru Miyoshi-Akiyama, Tomoko Sumitomo, Masanobu Nakata, Tadayoshi Ikebe, Tomoki Hanada, Takahiro Yamaguchi, Ryuji Kawahara, Rumi Okuno, Hitoshi Otsuka, Yuko Matsumoto, Yuji Terashima, Yu Kazawa, Noriko Nakanishi, Kaoru Uchida, Yumi Akiyama, Kaori Iwabuchi, Chikara Nakagawa, Kazunari Yamamoto, Shigetada Kawabata

## Abstract

Streptococcal toxic shock syndrome (STSS) caused by *Streptococcus pyogenes emm*89 strains has been increasing in several countries and reported to be linked with a recently emerged clade of *emm*89 strains, designated clade 3. In Japan, epidemiological and genetic information for *emm*89 strains remains elusive. In this study, we utilized *emm*89 strains isolated from both STSS (89 isolates) and non-STSS (72 isolates) infections in Japan from 2011 to 2019, and conducted whole-genome sequencing and comparative analysis, which resulted in classification of a large majority into clade 3 regardless of disease severity. In addition, STSS-associated genes and SNPs were found in clade 3 strains, including mutations of streptokinase (Ska), control of virulence sensor (CovS), serum opacity factor (SOF), sortase (SrtB), and fibronectin-binding protein F1 (PrtF1), and absence of the *hylP1* gene encoding hyaluronidase. These findings provide insights into notable genetic features of *emm*89 strains.

## Introduction

*Streptococcus pyogenes* is a human-specific pathogen known to cause a broad spectrum of diseases ranging from mild throat and skin infections to life-threatening invasive diseases (1, 2). Worldwide, it has been estimated that there are over 111 million cases of streptococcal pyoderma and 616 million cases of *S. pyogenes* pharyngitis, with 663,000 cases of invasive infection each year (3). The most severe manifestation noted is streptococcal toxic shock syndrome (STSS), which results in significant mortality with reported incidence rates ranging from 23% to 81% (4). Current estimates suggest that the incidence of STSS is increasing throughout the world, thus genetic characterization of recently emerged strains is useful for investigating novel therapeutic targets.

The M protein is a surface protein and one of *S. pyogenes* virulence factors (5). Although *S. pyogenes* typing has been historically conducted on the basis of M protein antigenicity, sequence typing of the *emm* region encoding the hyper-variable region of the M protein has also been widely applied as another method and used to classify the organism into at least 240 *emm* sequence types (6, 7). In recent years, increased incidence of STSS caused by *emm*89 *S. pyogenes* strains has been reported in both Europe and North America, with genome sequence analysis findings of *emm*89 strains used to identify 3 major genetically distinct strain clusters, designated as clade 1, 2 and 3 (8-10). Importantly, concurrent with the period in the early 2000s that featured a significant increase in number and frequency of invasive *emm*89 infections, clade 3 strains emerged and showed rapid expansion (8, 11).

Clade 3 strains are characterized by absence of the *hasABC* locus, which is responsible for synthesis of the hyaluronic acid capsule, as well as the presence of the *nga-ifs-slo* locus variant, associated with increased expressions of NADase and cytolysin (9, 11). Those reports indicated that most isolates from patients with STSS (STSS isolates) can be classified into clade 3. On the otherhand, recent distributions of *emm*89 clades in pharyngeal and asymptomatic isolates (non-STSS isolates) have not been thoroughly investigated. In Japan, the incidence of STSS caused by *emm*89 *S. pyogenes* is increasing (12), though the epidemiology of *emm*89 *S. pyogenes* strains, such as clade classification, remains unknown.

Whole genome sequence analysis of *S. pyogenes* strains can reveal polymorphisms among strains of the same *emm* type, such as classification of *emm*89 clades (8). One of the largest contributors to this genome-to-genome variation is small sequence differences, such as single nucleotide polymorphisms (SNPs). In the present study, we examined both STSS and non- STSS isolates of *emm*89 *S. pyogenes* in Japan obtained from 2011 to 2019, and conducted whole-genome sequencing and comparative analyses. The results classified most of the *emm*89 *S. pyogenes* collected in Japan as clade 3 regardless of disease severity. Furthermore, division of clade 3 strains into STSS- and non-STSS-associated phylogenetic groups was possible based on genome-wide SNP variations. We found both STSS-associated genes and SNPs in the clade 3 strains. These findings provide a fuller understanding of the recently emerged clade 3 of *emm*89 *S. pyogenes*, along with useful information for establishing novel strategies for treatment of diseases related to this pathogen.

## Materials and Methods

### Bacterial isolates

Clinical isolates were sent from multiple medical institutions to public health institutions. The diagnostic criteria used for determining *S. pyogenes*-induced STSS were based on definitive cases described by the Working Group on Severe Streptococcal Infections (1993) (13). The clinical isolates from STSS and non-STSS infections were collected during 2011-2019. STSS clinical isolates from sterile sites (89 isolates) and non-STSS clinical isolates from non-sterile sites were examined in this study (Supplementary, Supplementary Dataset 1). The number of STSS isolates collected in this study is estimated about 20-30% of the total number of incident cases reported. *emm* gene sequencing was performed as previously described by Beall *et al*. (14), with modifications as described at the Centers for Disease and Control website (https://www.cdc.gov/streplab/m-proteingene-typing.html).

### Ethical statement

The study protocol including use of human subjects was approved by the Ethics Committee of Osaka University Graduate School of Dentistry (Approval No: H29-E16-1). All procedures were performed in accordance with both ethical standards for human experimentation and the latest revised version of the Declaration of Helsinki.

### Genomic DNA sequencing

Isolates were cultured overnight in screw-capped glass tubes (Pyrex, Iwaki Glass. Tokyo, Japan) filled with Todd-Hewitt broth (BD Biosciences, San Jose, CA, USA) and supplemented with 0.2% yeast extract (BD Biosciences) (THY) at 37°C in an ambient atmosphere. Bacterial cells were lysed with 10 units/ml mutanolysin (Sigma, St. Louis, MO, USA), 10 mg/ml lysozyme (Wako, Osaka, Japan), and 0.5 mg/ml achromopeptidase (Wako), and genomic DNA was extracted from overnight cell cultures using a Maxwell^®^ RSC instrument with a Maxwell^®^ RSC PureFood GMO and Authentication Kit (Promega Corporation, Madison, WI, USA). Paired-end libraries were generated from extracted DNA with a Nextera^®^ XT DNA kit (Illumina, San Diego, CA, USA). Libraries were sequenced with Illumina instruments (HiSeq X and Miseq) and paired-end sequence reads (150 bp and 301 bp, respectively) were obtained.

### Data preprocessing and phylogenetic tree visualization

For subsequent genetic analyses, obtained sequence data were preprocessed to remove adapters and low-quality sequences with BBduk (https://github.com/BioInfoTools/BBMap). Next, obtained data were assembled *de novo* using a Unicycler (https://github.com/rrwick/Unicycler). Assembled contig and complete genome sequences (Supplementary Dataset 2) were analyzed using the kSNP3 program, version 3.1.1 (15), which is able to identify pan-genome SNPs in a set of genome sequences and estimate phylogenetic trees based upon those SNPs. The Kchooser program in kSNP3 was used to estimate the optimum k-mer values. Maximum likelihood trees were visualized using the Geneious Prime 2019.2 software package (Biomatters, Auckland, New Zealand).

### Detection of single nucleotide polymorphisms

Preprocessed reads were also aligned to the genome of MGAS23530, an *emm*89 clade 2 reference strain, using the Bowtie2package(http://bowtie-bio.sourceforge.net/bowtie2/index.shtml). Since the genome of that strain lacks prophages, and integrative and conjugative elements (8), the resultant polymorphism calls were made relative to only the core chromosome. Single nucleotide polymorphisms (SNPs) identified relative to the 1,709,394 nucleotides of MGAS23530 were concatenated and visualized using Geneious Prime 2019.2. A chi-square test was then used to determine associations between amino acid substitutions caused by SNPs and STSS.

### Screening of bacterial strains for hyaluronidase activity

After culturing overnight, 10 μL of *S. pyogenes* was inoculated onto the surface of a hyaluronan agar plate containing Todd-Hewitt broth (BD Biosciences), 1% yeast extract (BD Biosciences), 400 μg/mL hyaluronic acid sodium salt (Wako), 1% bovine serum albumin (Sigma), and 2% Noble agar (BD Difco, Franklin Lakes, NJ, USA). Following overnight incubation, agar plates were flooded with 2 N acetic acid solution and further incubated for 15 minutes. Strains exhibiting a halo zone around the colony were deemed positive for hyaluronidase activity. *S. pyogenes* strain 5448 (serotype M1, negative for hyaluronidase activity) and strain 4063-05 (serotype M4, positive for hyaluronidase activity) were employed as positive and negative controls, respectively (16).

### Data access

Data for the 161 sequenced *emm*89 genomes were deposited into the DDBJ sequence read archive (DRA) under accession number DRA009110.

## Results

### High-isolation rates of emm89 S. pyogenes clade 3

An important feature for characterization of clade 1, 2, and 3 is the *nga* promoter region sequence (Figure 1A). That sequence of *emm*89 clade 3 strains is identical to the *nga* promoter region present in pandemic *emm*1 strains (11). Furthermore, the *nga* promoter region sequence of clade 3 strains is associated with elevated production of *S. pyogenes* NADase and streptolysin O, secreted cytolytic toxins that contribute to its invasive phenotype (11, 17). Clade 3 *emm*89 strains also lack the *hasABC* region required for synthesis of the hyaluronic acid capsule (Figure 1B). We classified all *emm*89 strains into clade 1, 2, or 3 based on both the *nga* promoter region sequence and presence of *hasABC* genes. Regardless of disease severity, nearly all (96.9%) of the strains were determined to be clade 3 (Supplementary Dataset 1), while only 5 showed characteristics of clade 2 strains and no clade 1 strain was detected. Chi-square test results indicated that clade 3 is not associated with STSS (Table 1).

**Table 1.** Factors associated with STSS.

| Factor | RefSeq locus tag | Gene<br>symbol | STSS<br>(n=89) | Non-STSS<br>(n=72) | $\chi^2$ value* | P-value |
| --- | --- | --- | --- | --- | --- | --- |
| Associated with STSS |  |  |  |  |  |  |
| Mutation in CovS | MGAS23530_0304 | *covS* | 20 | 2 | 13.29 | 2.67.E-04 |
| Absence of *hylP1* gene | MGAS27061_0574 | *hylP1* | 34 | 10 | 11.85 | 1.00.E-03 |
| Not associated with STSS |  |  |  |  |  |  |
| Clade 3 |  |  | 87 | 69 | 0.49 | 4.85.E-01 |
| Mutation in CovR | MGAS23530_0303 | *covR* | 4 | 0 | 3.32 | 6.85.E-02 |
| Mutation in Rgg | MGAS23530_1555 | *rgg* | 3 | 4 | 0.46 | 4.99.E-01 |

\* $\chi^2$ values for evaluating association between STSS and factors.

**Figure 1.**
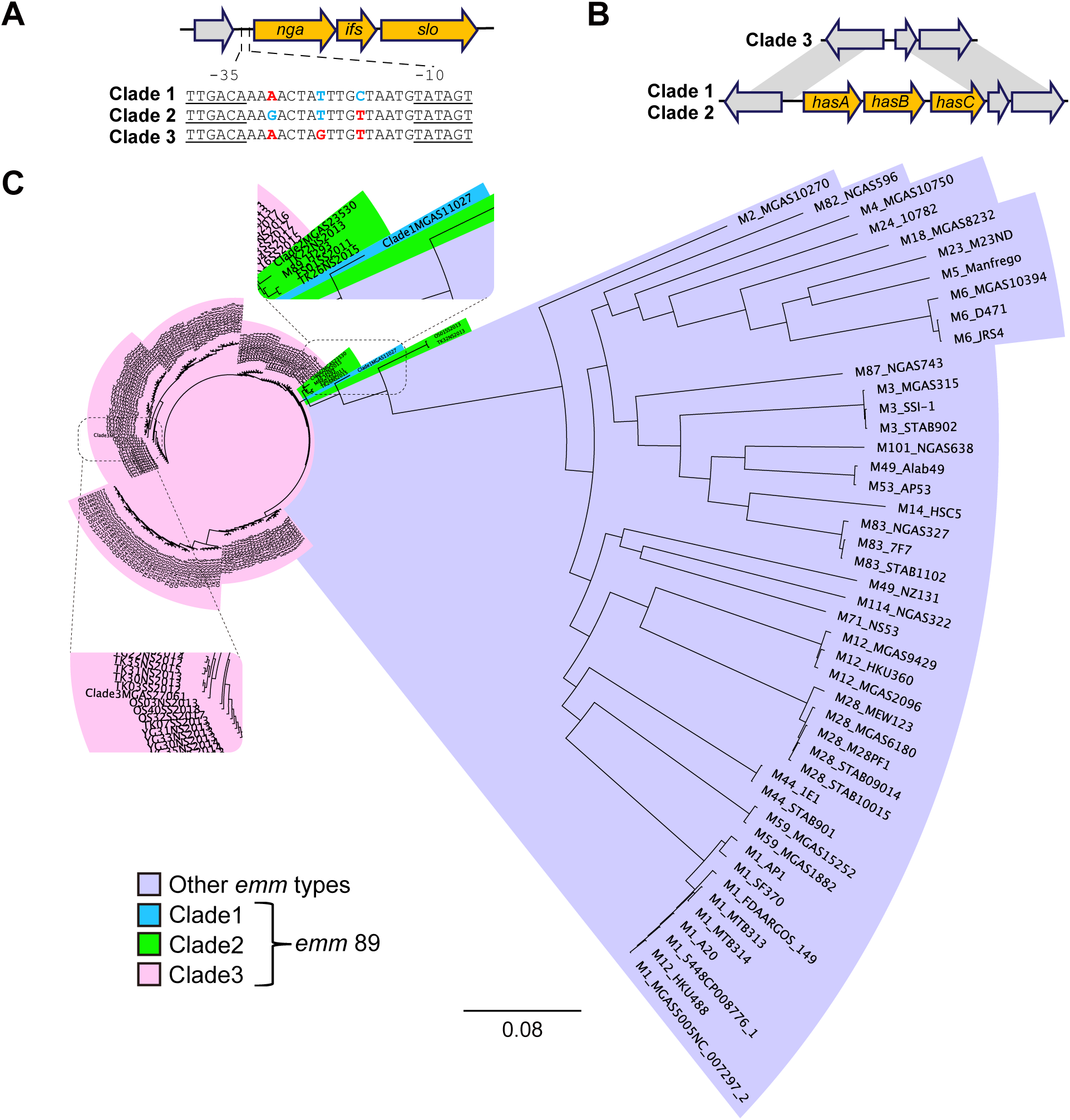
Classification criteria for clade, and genetic relationships among various *S. pyogenes* strains. (A) Schematic diagram showing *nga* promoter region sequences for each clade. (B) Schematic alignment of *hasABC* operon for clade 1, 2, and 3. *hasABC* was not found in clade 3. (C) Genetic relationships between *emm*89 strains in Japan and various *emm*/M protein serotype strains. Genetic relationships were inferred among 49 GAS strains of 23 *emm*/M types based on 73,030 concatenated core chromosomal SNPs using the maximum likelihood method. Three closed genomes (clade 1, MGAS11027; clade 2, MGAS23530; clade 3, MGAS27061) were used as references. Scale bar indicates nucleotide substitutions per site. *S. pyogenes emm*89 strain clade 1, 2, and 3 are shaded in blue, green, and pink, respectively. Various *emm*/M protein serotypes except *emm*89 strains are shaded in purple. FS, regions containing Sapporo city, Iwate prefecture, Fukushima prefecture, Sendai city, and Niigata city; TY, Toyama prefecture; TK, Tokyo prefecture; YH, Yokohama city; OS, regions containing Shiga prefecture, Kyoto city, Osaka prefecture, and Hyogo prefecture. YG, Yamaguchi prefecture. SS, isolates from STSS patients; NS, pharyngeal or asymptomatic isolates. 20xx indicates year of isolation.

### STSS- and non-STSS-associated phylogenetic groups among emm89 clade 3 strains

We constructed kSNP trees using 161 *emm*89 assembled contigs and 49 other *S. pyogenes* complete genome sequences (Figure 1C) (for information regarding strains used in this study see Supplementary Dataset 1 and 3). As control genomes of the *emm8*9 strains, 3 reference genomes from representative pre-epidemic (clade 1, MGAS11027; clade 2, MGAS23530) and epidemic (clade 3, MGAS27061) *emm*89 strains sequenced in a previous study by Zhu *et al*. were used (18). The phylogenetic tree indicated a genetically close relationship of the 161 *emm*89 *S. pyogenes* strains isolated in Japan. The 161 *emm*89 strains were clustered with the reference genome of clade 2 (MGAS23530) or clade 3 (MGAS27061), and the cluster groups were consistent with the results of classification of clades based on both *nga* promoter region sequences and presence of *hasABC* genes (Supplementary Dataset 1).

The *S. pyogenes emm*89 strains in the phylogenetic tree show a close evolutionary relationship and it is difficult to follow their branching relationship (Figure 1C). Therefore, we drew a cladogram to more clearly show branching relationships among the strains (Supplementary Figure 2). In addition, we constructed kSNP trees using assembled contigs of the 161 *emm*89 strains and 3 control genomes (Figure 2, 3, Supplementary Figure 3, 4). There was no regularity in regard to regional (Figure 2A) or temporal distribution (Supplementary Figure 4). On the other hand, we found that strains classified as clade 3 could be divided into STSS- and non-STSS-associated phylogenetic groups, which were termed the SS and NS groups, respectively (Figure 2B, 3).

**Figure 2.**
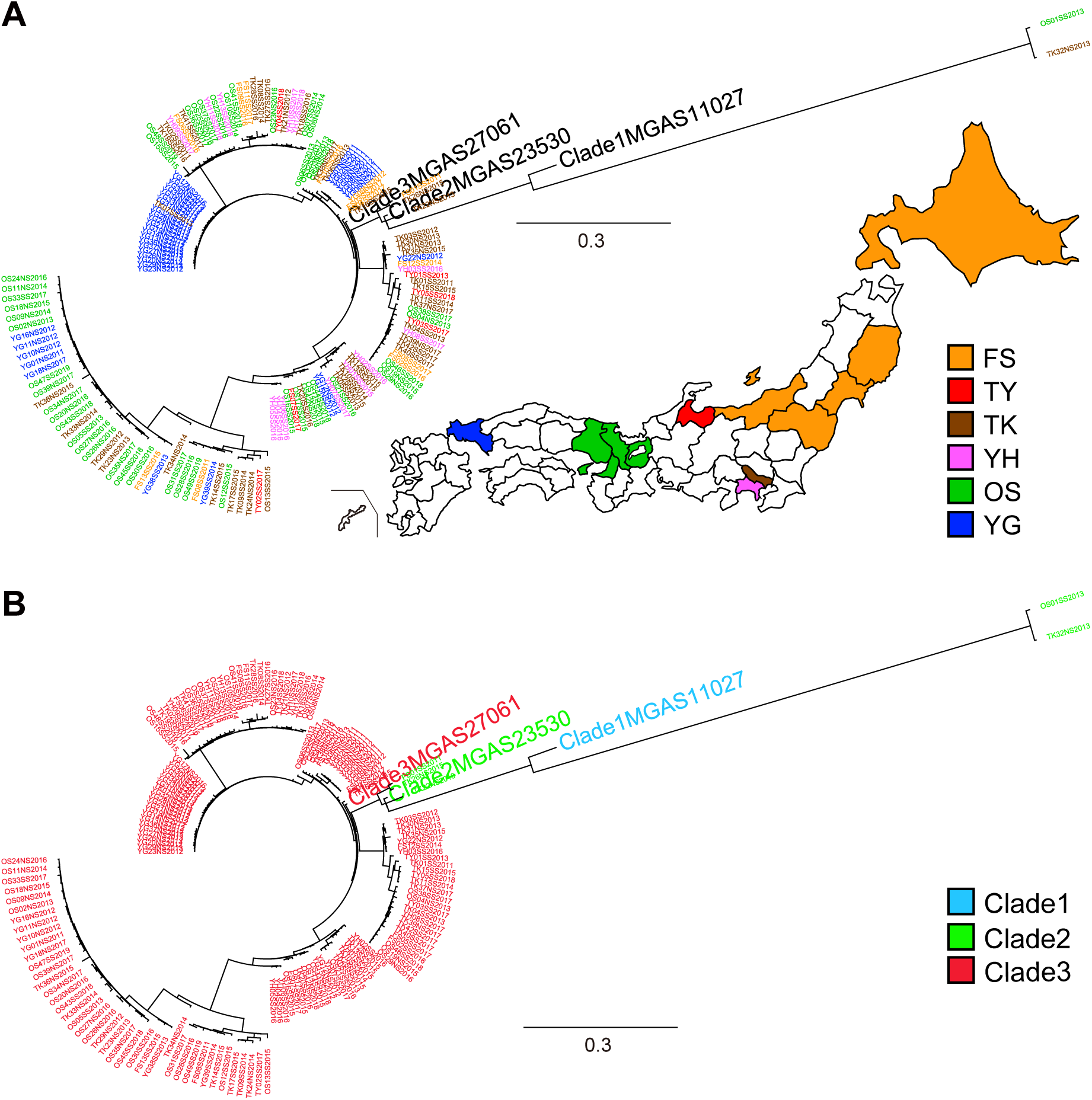
Regional and clade distributions of *emm*89 strains in Japan. (A) Regional distribution of *emm*89 strains in Japan. Genetic relationships between *emm*89 strains in Japan and *emm*89 reference strains (clade 1, MGAS11027; clade 2, MGAS23530; clade 3, MGAS27061) are shown. Genetic relationships were inferred among 164 strains based on 8866 concatenated core chromosomal SNPs using the maximum likelihood method. Strains are colored based on region of isolation, as indicated in the index. (B) Clade distribution of *emm*89 strains in Japan. Strains are colored based on clade, as indicated in the index.

**Figure 3.**
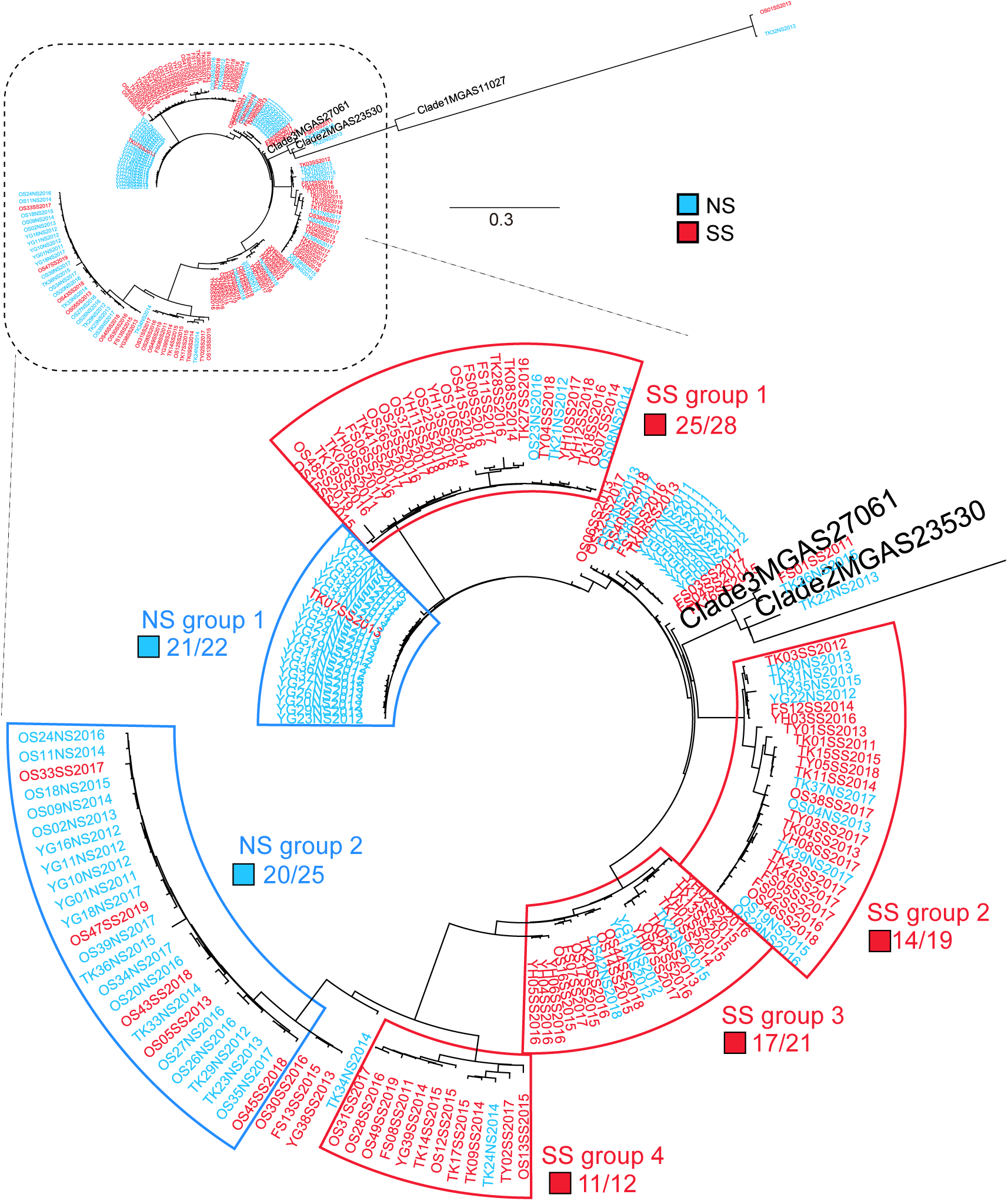
STSS- and non-STSS-associated phylogenetic groups among *emm*89 clade 3 strains. Strains are colored based on SS (STSS isolates) or NS (non-STSS isolates), as indicated in the index. Strains are classified as STSS- or non-STSS-associated phylogenetic groups, and termed SS (SS group 1, 2, 3, 4) and NS (NS group 1, 2) groups, respectively.

**Figure 4.**
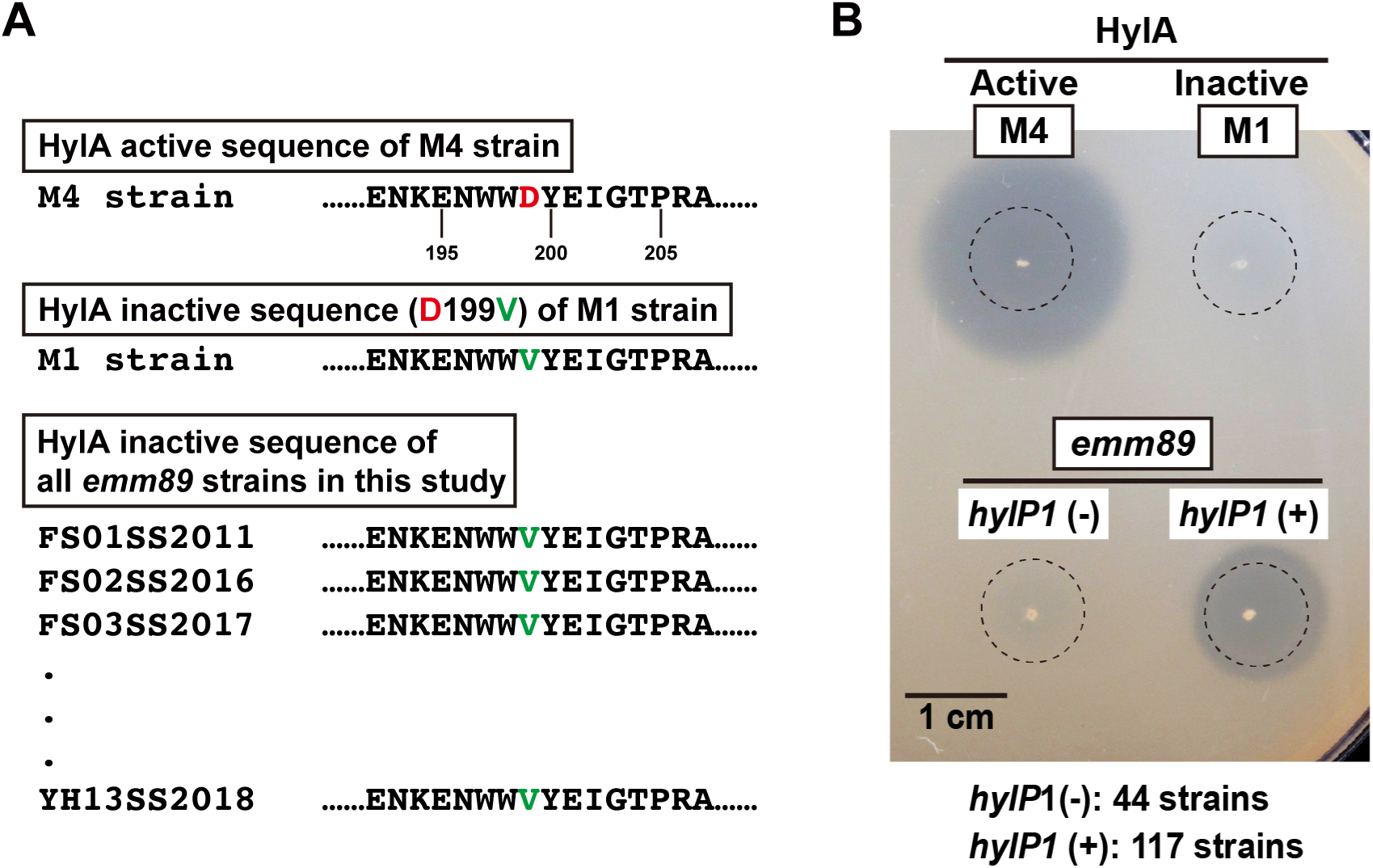
Hyaluronidase activity of *emm*89 *S. pyogenes*. (A) Hyaluronidase HylA active or inactive sequence. Serotype M4 strain was shown to harbor HylA activity and serotype M1 strain possession of Asp to Val substitution at position 199 of the HylA sequence (D199V), which abolishes hyaluronidase activity (16). (B) Hyaluronidase activity on hyaluronan agar plate. Dotted circles indicate colony locations before incubating with 2 N acetic acid solution. Although 161 *emm*89 strains in this study harbored D199V in the HylA sequence, 117 of the strains were indicated to have hyaluronidase activity.

### Amino acid mutations contributing to division of groups

Preprocessed reads of the 161 *emm*89 strains were aligned to the genome of the *emm*89 clade 2 reference strain MGAS23530, which revealed a polymorphism involved in the division of groups (Supplementary Dataset 3) and associated with STSS (Table 2, Supplementary Dataset 3). Regarding virulence factors, an amino acid substitution of streptokinase (Ska, Ile to Thr substitution at position 17: I17T) was associated with STSS, while amino acid substitutions of NAD glycohydrolase (Nga, D130G) and streptolysin S biosynthesis protein C (SagC, P223S) were associated with non-STSS. As for factors with effects on cell surface molecules, amino acid substitutions of fibronectin-binding protein F1 (PrtF1, F420L), sortase (SrtB, V131A), and serum opacity factor (SOF, G843S, M844V) were found to be associated with STSS, and those of collagen-like surface protein (SclB, P180A and A228P) was found to be associated with non- STSS. Additionally, an amino acid substitution of ferrichrome transport system permease protein (FhuB, V73A) was associated with non-STSS. These results suggest that specific SNPs contribute to the invasiveness of individual clade 3 strains.

**Table 2.**
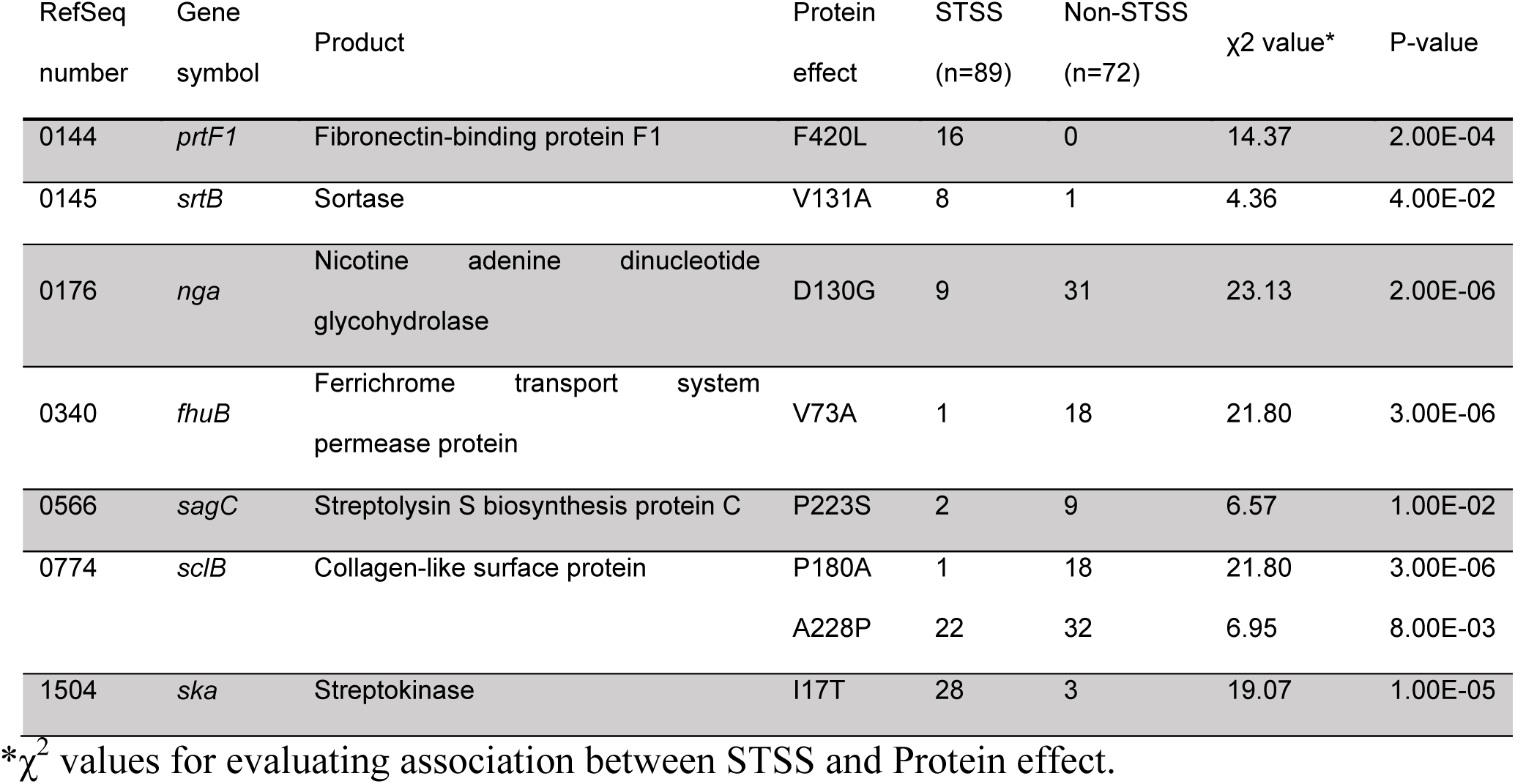
Phylogenetic group-associated mutations and association with STSS.

### Other STSS-associated factors

Mutations of the CovRS virulence regulator derepress virulence genes, leading to a hypervirulent phenotype of *S. pyogenes* (19, 20). Of the 161 clinical isolates of *emm*89 *S. pyogenes* identified in Japan, 20 STSS isolates (22.5%) and 2 non-STSS isolates (2.7%) showed CovS mutations (p = 0.000267, χ^2^ analysis) (Table 1, Supplementary Dataset 1). On the other hand, the frequency of CovR mutations in the isolates was not associated with STSS. Although the Rgg mutation has also been reported as an important factor in the pathogenesis of invasive infections (21), that was not found to be relevant to STSS in this study.

Hyaluronidase secreted by Group B *Streptococcus* cleaves pro-inflammatory hyaluronan, which contributes to evasion of host immunity (22). Accordingly, we screened the 161 isolated *emm*89 strains for hyaluronidase activity to investigate their involvement in STSS. Serotype M4 strain 4063-05 was used as a positive control for hyaluronidase (HylA) activity, while the negative control was serotype M1 strain 5448, which possesses an Asp to Val substitution at position 199 of the HylA sequence (D199V) that abolishes hyaluronidase activity (16). Although all 161 strains harbored D199V in the HylA sequence, 117 showed hyaluronidase activity when cultured on hyaluronan agar plates (Figure 4, Supplementary Dataset 1), and that activity was consistent with the presence of the *hylP1* gene (Supplementary Dataset 1). The *hylP1* gene was absent in 34 (38.2%) of the STSS isolates and 10 (12.7%) of the non-STSS isolates (p = 0.001, χ^2^ analysis) (Table 1). HylP1 is one of the key bacteriophage-encoded virulence factors (23). However, in regards to S. *pyogenes emm*89, absence of *hylP1* gene is relevant to STSS.

## Discussion

Our results indicate that most *emm*89 strains isolated in Japan since 2011 are members of the recently emerged genetic clade 3, regardless of disease severity. Similar findings have been reported in Europe, including Finland, Iceland, England, and Portugal, as well as North America (United States and Canada) where clade 3 *emm*89 strains have shown recent emergence. Reports from countries other than Japan suggest that recently documented invasive infections caused by *emm*89 strains are due to the appearance of clade3. However, in the present study based in Japan, we found genes and SNPs associated with STSS as well as non-STSS in clade 3 strains. Some SNPs in virulence factors and adhesins were detected.

Streptokinase (Ska) is highly specific for human plasminogen and contributes to streptococcal virulence by generating plasmin, which leads to bacterial spread from a primary focus of infection by causing fibrinolysis, as well as degradation of the extracellular matrix and basement membrane components (24). The Ile or Thr residues at position 17 of the Ska sequence of *emm*89 strains are located in the streptokinase α domain. The α domain contributes to interactions between the catalytic domain of human plasminogen and streptokinase (25). A substitution at position 17 of the Ska sequence (I17T) may enhance streptococcal virulence by strengthening enzyme activity.

NADase and streptolysin S are virulence factors involved in disease pathogenesis associated with *S. pyogenes* (17, 26). Although no studies have reported a Pro to Ser substitution at position 223 of the SagC sequence (P223S), nor an Asp to Gly substitution at position 130 of the Nga sequence (D130G), those mutations may be associated with decreased enzyme activity because of their association with non-STSS isolates.

The *prtF1* and *srtB* genes are located in the FCT (<u>F</u>ibronectin and <u>C</u>ollagen-binding proteins and <u>T</u>-antigen) region, where pilus genes are also located (27, 28). Fibronectin-binding protein F1 (PrtF1) of *S pyogenes* has roles in both adherence to and internalization in respiratory epithelial cells (29). Although SrtB was found to be primarily required for efficient formation of the backbone protein T6 and FctX complex, and subsequent polymerization of T6 in a serotype M6 strain (classified as FCT type 1) (30), it remains unknown whether SrtB of *emm*89 strains (classified as FCT type 4) contributes to pilus formation. A Phe to Leu substitution at position 420 of the PrtF1 sequence (F420L) and a Val to Ala substitution at position 131 of the SrtB sequence (V131A) were shown to be relevant in the present STSS isolates, and those substitutions may enhance the virulence of *S. pyogenes* by changing the functions of PrtF1 and SrtB. However, note that the effects of these SNPs on molecular functions were not identified, and further verification is necessary to prove the relevance to STSS.

Ikebe *et al*. reported that CovRS and Rgg mutations have important roles in the pathogenesis of STSS (21). The present results showed mutations of CovS associated with STSS, while those of CovR and Rgg did not have such an association. However, it is possible that relevance could not be determined due to the small number of strains examined.

*emm*89 clade 3 strains lack hyaluronic acid capsule synthesis genes (*hasABC*), as previously reported for the *S. pyogenes* M4 and M22 strains (31). M4 and M22 are the only *S. pyogenes* strains known to possess an active HylA sequence (16), whereas that was not found in the *emm*89 strains examined in the present study. Therefore, a genomic type of *emm*89 clade 3 strain harboring an inactive HylA sequence, even though *hasABC* is deleted, has yet to be reported. We also found no association of STSS with HylP1-inactive *emm*89 strains. It was previously reported that an HylA-deficient mutant of *S. pyogenes* M4 did not display a significant reduction in virulence as compared with a wild-type strain (32). Together, these findings suggest that hyaluronidase activity may be dispensable for invasive infection by a specific subset of *S. pyogenes* strains.

*S. pyogenes emm*89 clade 3 strains have recently emerged in Japan. The present whole-genome sequence results provide some important genetic information for additional research to characterize genetic variations of *S. pyogenes* as well as findings showing novel notable features related to molecular epidemiology. Knowledge obtained in this study may also help to provide a more detailed molecular characterization as well as molecular targets for developing treatment strategies for patients with diseases related to *S. pyogenes* infection.

## Acknowledgments

This study was supported in part by AMED (JP19fk0108044, JP19fm0208007), Japanese Society for the Promotion of Science (JSPS) KAKENHI (grant numbers 19H03825, 20K18474), JSPS Overseas Research Fellowships, Takeda Science Foundation, Japanese Association for Oral Biology Grant in Aid for Young Scientists, SECOM Science and Technology Foundation, The Naito Foundation, and Kobayashi International Scholarship Foundation. The funders had no role in study design, data collection or analysis, decision to publish, or preparation of the manuscript.

The authors wish to express their gratefulness to Y. Nakamura (Sapporo City Institute of Public Health) and N. Hashimoto (Sendai Municipal Institute of Public Health), as well as the medical institutions that participated in collection of the present strains.

